# Computational Analysis of Dynamic Allostery and Control in the three SARS-CoV-2 non-structural proteins

**DOI:** 10.1101/2020.12.12.422477

**Authors:** Igors Dubanevics, Charles Heaton, Carlos Riechmann, Tom C.B. McLeish

## Abstract

Severe acute respiratory syndrome coronavirus 2 (SARS-CoV-2), which caused the COVID-19 pandemic, has no vaccine or antiviral drugs available to the public, at the time of writing. The virus’ non-structural proteins are promising drug targets because of their vital role in the viral cycle. A significant body of work has been focused on finding inhibitors which covalently and competitively bind the active site of the non-structural proteins, but little has been done to address regions other than the active site, i.e. for non-competitive inhibition. Here we extend previous work on the SARS-CoV-2 M^pro^ (nsp5) to three other SARS-CoV-2 proteins: host shutoff factor (nsp1), papain-like protease (nsp3, also known as PLpro) and RNA-dependent RNA-polymerase (nsp12, also known as RdRp) in complex with nsp7 and nsp8 cofactors. Using open-source software (DDPT) to construct Elastic Network Models (ENM) of the chosen proteins we analyse their fluctuation dynamics and thermodynamics, as well as using this protein family to study convergence and robustness of the ENM. Exhaustive 2-point mutational scans of the ENM and their effect on fluctuation free energies suggest several new candidate regions, distant from the active site, for control of the proteins’ function, which may assist the drug development based on the current small molecule binding screens. The results also provide new insights, including non-additive effects of double-mutation or inhibition, into the active biophysical research field of protein fluctuation allostery and its underpinning dynamical structure.

## 1. Introduction

During 2020, a rapidly spreading viral disease, COVID-19, caused by the novel coronavirus SARS-CoV-2, has generated a global pandemic. Although early fast-tracked vaccines are in development, no vaccine or specific anti-viral drugs are publicly available at the time of writing. Furthermore, in the longer term, the identification of all potential inhibitor sites at all points of the viral life-cycle is of interest in support of a flexible pharmaceutical response. In this work we focus on the low-frequency dynamical structure of three the virus’ proteins important for the viral cycle. These are host shutoff factor (nsp1), papain-like protease (nsp3, also known as PLpro) and RNA-dependent RNA-polymerase (nsp12, also known as RdRp) in complex with nsp7 and nsp8 cofactors. To this end we develop and test an elastic network model (ENM) - based methodology recently tested on the SARS-CoV-2 M^pro^ (1) that analyses the low-frequency structure of protein dynamics. Since the ENM analysis permits rapid scanning of potential binding and mutation sites that offer critical control of the protein dynamics, we identify critical residues for potential allosteric control of their function. The work is informative for inhibitor design by identifying control regions of the proteins that are distant from, rather than proximal to, its active sites. Computational studies such as this one may go hand-in-hand with rapid experimental binding scans of small-molecules (2, 3), to identify those agents most likely to inhibit protein function as those which bind to sites critical to the functional dynamics.

Allosteric mechanisms for distant control of binding and activation fall into two main classes: those which invoke significant conformational change (the original scenario of Monod, Wyman and Changeaux (4), and mechanisms that invoke the modification of thermal (entropic) fluctuations about a fixed, mean conformation (5–8). Such ‘fluctuation allostery’ recruits mostly global, low-frequency modes of internal protein motion, which are well-captured by correspondingly coarse-grained mechanical representations of the protein (9, 10). One effective tool at this level is the Elastic Network Model (ENM) (11). The ENM resolves protein structure at the level of alpha-carbon sites only, which are represented as nodes connected by harmonic springs within a fixed cut-off radius from each other. Local point mutation can be modelled by changing the moduli of springs attached to the corresponding residue, and effector-binding by the addition of nodes (and local harmonic potentials) at the corresponding co-ordinates. The most significant contributions to both the correlated dynamics of distant residues, and to the entropy arising from structural fluctuation, come from global (’low frequency’) modes, which are well-captured by the ENM approximation. This approach was successfully used to identify candidate control residues whose mutation may control allostery of effector-binding in the homodimer transcription factor CAP (12), and very recently to identify similar candidates for functional control of the SARS-CoV-2 M^pro^ (1), some of which have been identified experimentally.

These studies have shown that, while the ENM approximation (ignoring e.g., side-chain structure) does not capture the quantitative values of free energies, it does rank their values qualitatively, and correctly identifies the functional form of the protein’s low-frequency modes, as well as residues which present as candidates for allosteric control through mutation. The method, and the open software (’DDPT’) used in the previous studies on allosteric homodimers, and confirmed by experimental calorimetry on model-designed mutations (13), is deployed here in a similar way (see Methods section) to a coarse-grained ENM models of three further SARS-CoV-2 proteins.

There are a number of free parameter choices and possible extensions of the ENM that present themselves as candidates for optimising the reliability and robustness of its results on dynamic protein analysis. In particular, the choice of cut-off distance below which the ENM model constructs a harmonic potential between any two C-alpha atoms of a protein, is a parameter that needs to be chosen with care. Too small a cut-off risks creating a model with sub-critical connectivity and containing unphysical normal modes of zero elastic modulus. Too large a cut-off creates an over-rigid model of the protein, which fails to capture important low-frequency global modes of motion. Ideally one might expect a universal optimum value for the cut-off, but the question arises of legitimate protein-dependent tuning. Secondly, the simplest ENM model imposes a universal bond stiffness, irrespective of whether two C-alpha carbons are neighbours along the protein backbone, or proximate because of the native state fold. The covalent forces in the first case would be expected to deliver stronger stiffness than the non-covalent bonds in the second. An extension to the ENM that contains two populations of spring constants corresponding to these two populations was investigated by Ming and Wall (14). In the light of these aspects of residual freedom within the set of ENM models, we exploit this set of studies of SARS-CoV-2 proteins to examine the effect of such enhanced ENMs, and of varying cut-offs in bond distance and normal mode sum on the key predictions of allosteric control, in order to improve an understanding of the ENM coarse-graining process.

In the following, section 2 summarises the results on SARS-CoV-2 M^pro^ reported in (1), extending them to explore the reliability and robustness of the ENM approach in terms of (i) cut-off distance, (ii) cut-off in mode number, (iii) choice of backbone stiffness constant. Then section 3 presents the results of ENM analysis on the host shutoff factor (nsp1), the papain-like protease (nsp3) and RNA-dependent RNA-polymerase (nsp12). The final, concluding, section reviews the potential for small-molecule inhibition of these proteins, and the additional physics of protein dynamics that they serve to expose.

## 2. The SARS-CoV-2 Main Protease Protein: Review and ENM Model Development

In its active form M^pro^ is a two-protomer homodimer with one active site per homodimer chain (15). A recent study characterised a SARS-CoV M^pro^ mutation, S284-T285-I286/A, which dynamically enhanced the protease catalytic activity more than three-fold (16).

### A. ENM Results for Allosteric Control

Such experimental findings support the hypothesis of dynamically driven allosteric control of SARS-CoV-2 M^pro^, and provide a structure (6lu7) on which to base an ENM construction. All three PDB files used in this study were derived from original crystallographic structure of SARS-CoV-2 M^pro^ with N3 inhibitor.

A recent crystal structure for the SARS-CoV-2 M^pro^ is shown in front view in figure 1, alongside the ENM model of (1), and the structure of the first non-trivial normal mode. That work took C_*α*_ node masses as the whole residue mass, and uses a cut-off distance for harmonic connecting springs of 8Å, based on optimising the comparison of mode structures between ENM and full Molecular Dynamics simulations, in previous work on Catabolite Activator Protein (12, 17). Away from the dimer interface, the protein structure is dominated by beta-sheets, while the interface itself is composed of adjacent helices and loops. Although M^pro^ is a relatively compact protein (fewer than 310 residues per chain), it plays a vital role in the viral cycle of both coronaviruses: it divides polyproteins expressed from the viral mRNA into its functional non-structural units (18). This functional role makes SARS-CoV-2 M^pro^ an appealing target for drug design.

**Fig. 1.**
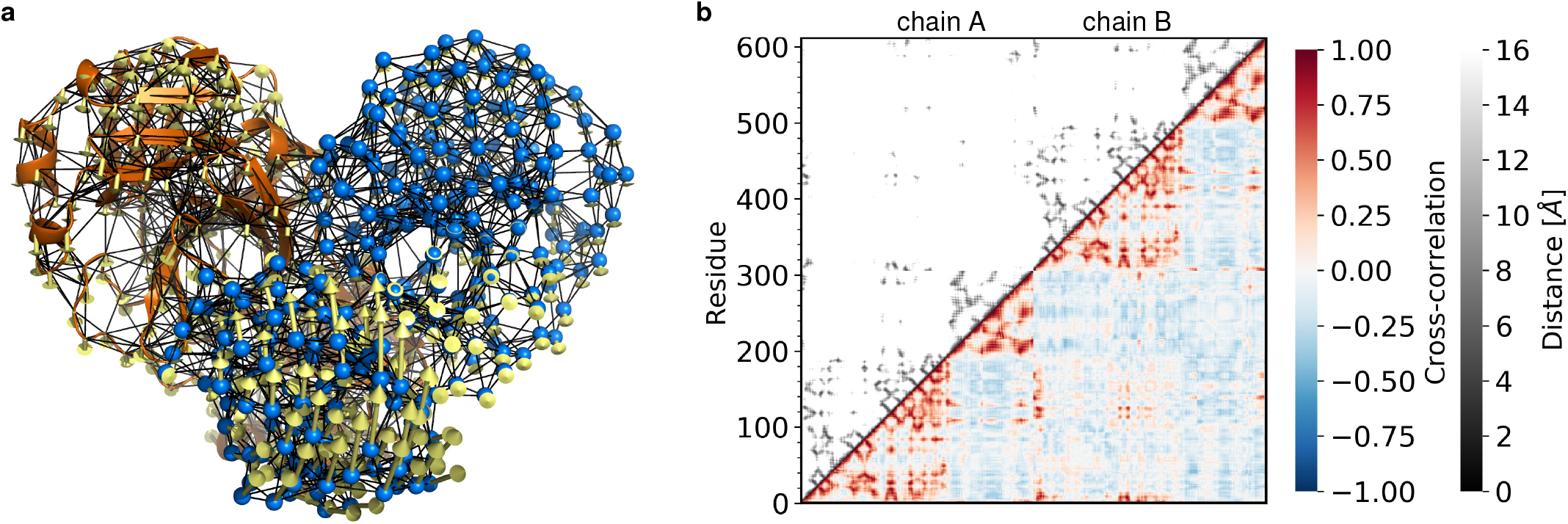
Dynamics of the SARS-CoV-2 M^pro^ ENM. **a**, Secondary structure cartoon (orange); C_*α*_ atom nodes (marine); node-connecting springs in black; the first real fluctuation mode eigenvector displacement (yellow) are scaled 5 times. The ENM was generated with PyANM plugin in PyMOL. **b**, The cross correlation of the motion of 6lu7 apo ENM. The cross-correlation maps calculated for the first real 25 modes (bottom-right region of the plots) and spacing between residues (C_*α*_ nodes) and ligand nodes (top-left region of the plots) for the apo form. The first colour scale shows the extent of cross correlation, with a cross correlation of 1 (red) indicating perfectly correlated motion, −1 (blue) showing perfectly anti-correlated motion and 0 (white) no correlation. The second colour scale (black to white) depicts the Euclidean distance between two ENM’s nodes in the Cartesian space in 0-16 Å range.

In case of the bound N3 inhibitor, the short polypeptide AVL was coarse-grained in the same way as the main chain amino acids while the other heavy atoms (C, N and O) have been treated as individual nodes with atomic masses. The analysis showed that SARS-CoV-2 M^pro^ possesses a rich dynamical structure that supports several long-distance allosteric effects through thermal excitation of global normal modes. In particular the motions in the vicinity of two active sites are correlated within the first 25 non-trivial normal modes, especially in the singly-bound dimer. This correlation appears in the two-point dynamic correlation map derived from the ENM for the apo protein shown in figure 1b (lower half). The figure also displays a corresponding two-point distance map (top quadrant), from which it can be seen that most, but crucially not all, dynamic correlations arise through proximity. Although, at the level of ENM calculations, this does not lead to cooperativity in the WT structure, it does render the protein susceptible to the introduction of cooperativity by mutation.

Our methodology is further supported by the ENM dynamics sensitivity to residue 214 and 284-286 mutations which have been shown by experiment to dynamically control SARS-CoV M^pro^.The ENM calculations have identified new sites whose local thermal dynamics dynamically correlate with those of the active sites, and which also appear on global maps for allosteric control by single or double mutations. Examples of the reports available from the ENM calculations that expose such structure are given in figure 1, showing a dynamical residue-residue correlation map. Double-mutation scans are also instructive, and were presented in (1).

### B. Using the SARS-CoV-2 M^pro^ for ENM Model Development

The computations in (1) chose mode and distance cutoffs by balancing the requirements of: (i) sufficient spatial resolution of dynamics; (ii) requirements not to include unphysically small-scale structure; (iii) acceptable convergence of thermodynamic calculations; (vi) compatibility with the previous studies (19). These led to the choice of summing the first real 25 modes in SARS-CoV-2 M^pro^ ENM calculations.

Other previous work has suggested a simple development of the ENM model in which the different chemical character of main-chain and other bonds in globular proteins is recognised by allocating two, rather than one single, spring constant to ENM models (14). Specifically, main chain bonds are modelled by a stronger harmonic potential than all other bonds within the ENM cut-off, which continue with an otherwise uniform spring stiffness. In the light of the choice of parameters offered by the two cut-offs required in ENM models (bond distance and mode number), as well as the possibility of more accurate models using two spring stiffnesses rather than one, before embarking on the analysis of other key proteins in the SARS-CoV-2 family, we explored the parameter choice using the SARS-CoV-2 M^pro^ already explored in depth. Moreover, in this study we used a ENM with all elastic network node mass set to 1 atm. This simplification has been used in previous studies (12, 17, 19) and has shown no difference between the mass-weighted model when computing relative fluctuation free-energy change.

#### B.1. ENM Cut-offs in bond distance and normal mode sum

Figure 2a reports the average correlation of ENM and experimental b-factors over the entire protein structure for the SARS-CoV-2 M^pro^ homodimer as a function of the upper cut-off in the sum of normal modes, and for a range of cut-off distances used in assembling the ENM. The results agree with earlier studies that indicate that the bond-distance cut-off needs to be at the least 8Å in order to capture the experimental residue dynamics with acceptable accuracy. By the point at which the first 12 modes have been included in the sum, the model/experiment correlation is above 0.5. Adding higher modes only marginally improves agreement up to the first 100 modes, beyond which there is certainly unphysical structure being included. Beyond a bond distance cut-off of 8Å the correlation remains optimised with this behaviour up to a cut-off of 10Å, at which point there appears a small decrease in correlation as more modes are incorporated up to the 20th.

**Fig. 2.**
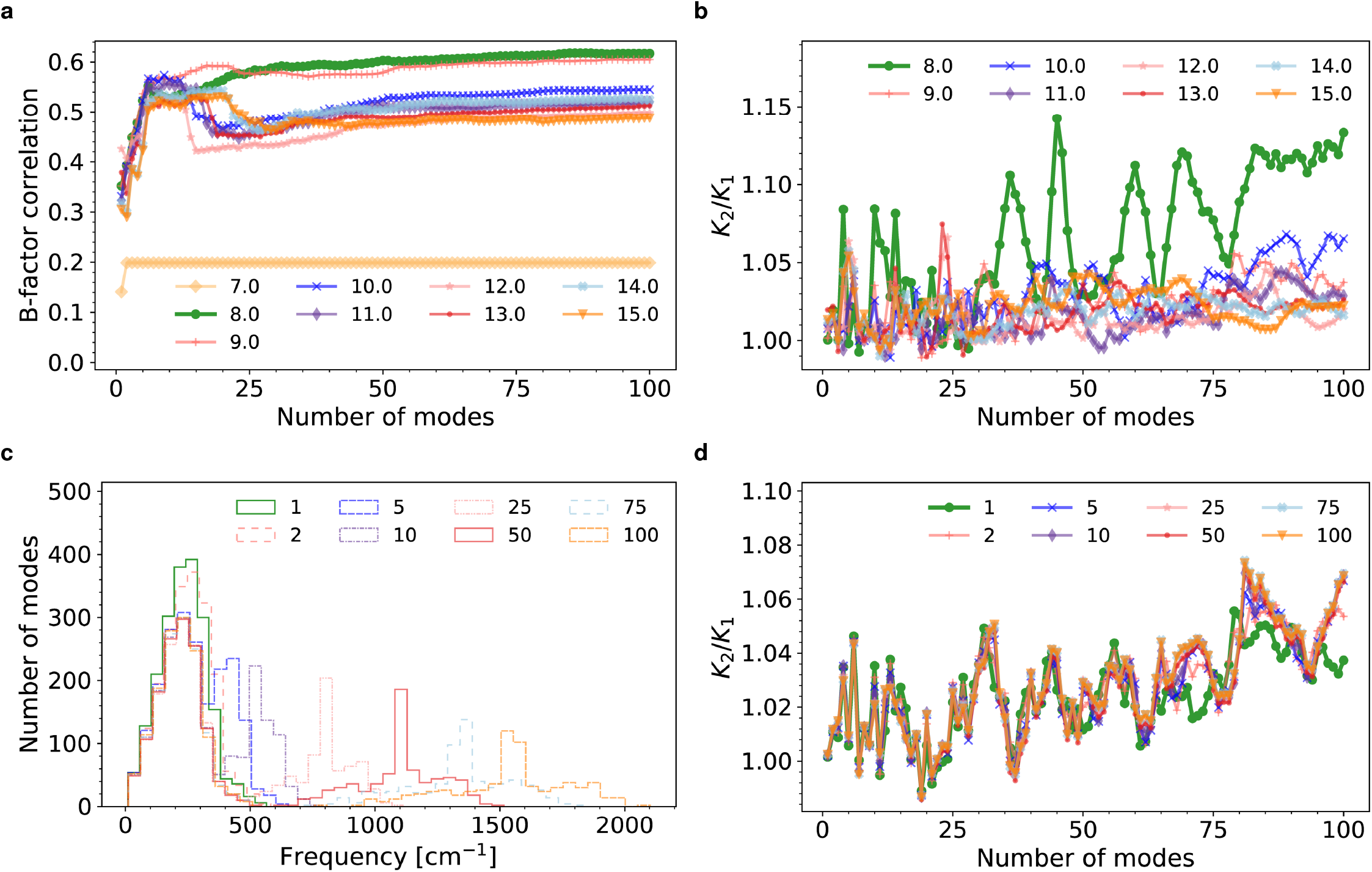
SARS-CoV-2 M^pro^ ENM response to the cut-off distance and backbone spring constant variation.**a**, B-factor correlation with the ENM normal modes with the real mode summation for a range of cut-off distances between 7 and 15 Å. **b**, Ligand dissociation ratio versus the mode summation for the cut-off in range of 8 to 15 Å. **c**, Density-of-states distribution for the ENM with varied backbone spring constant between 1 to 100 for 9Å cut-off. **d** Ligand dissociation ratio versus the mode summation for the backbone spring constant between 1 to 100 for 9Å cut-off.

The b-factor calculations were complemented, as a function of the same varying parameters, by comparing predictions of the allosteric free-energy. The allosteric measure used is the ratio of dissociation constants for second and first inhibitor binding *K*2*/K*1. The results, shown in figure 2b, are noisier than the b-factor computations. This is expected, since the allosteric free-energy is a second order difference of free energies, while the b-factor is a first-order quantity. Nevertheless, this measure displays a greater sensitivity to the lower limit for acceptable bond distance cutoff than does the b-factor, for in this case acceptable convergence is found only at a 9 Å cutoff and above. In regard to the mode-sum, convergence is optimised from 10-25 modes, after which adding further modes is visible, by comparing results with different distance cutoffs, as a noisier process. In conclusion, the two measures taken together suggest, for the standard ENM model, that a 9 Å - 10 Å cutoff together with a mode sum to 25 non-trivial modes is an optimal choice.

#### B.2. BENM (Backbone-enhanced Elastic Network Model)

Figure 2c reports the distribution function for normal mode stiffness for SARS-CoV-2 M^pro^ homodimer for a range of relative values of the main-chain stiffness, employing the previously-optimised choice of 9 Å for the bond distance cut-off. We find, just as did Ming and Wall in (14), that increasing the backbone stiffness generates a bimodal distribution of mode stiffness, rather than the single-peaked distribution of the standard ENM. From this point of view, the value of backbone stiffness appears crucial for a physically accurate coarse-grained model of protein dynamics.

One way to motivate a choice of backbone stiffness among the range of possible values, is to optimise the faithfulness of the ENM coarse-grained model to a finer-grained fully-atomistic model. To this end, an all-atom molecular dynamics simulation of the apo SARS-CoV-2 M^pro^ homodimer was made, using the AMBER potential scheme (20). After equilibration, data from a run of 20 ns simulated dynamics was used to identify the normal modes of motion of the set of C_*α*_ atoms (see methods section). Their distribution is also recorded in figure 2c, and like the ENM models with enhanced backbone stiffness, displays a bimodal distribution. However, none of the ENM models was able to capture both the frequency ratio of the two peaks and also their relative peak-integrals (intensities). Choosing to optimise the frequency ratio (of 4 seen in the AMBER simulations) suggests a ratio of backbone to non-backbone bond stiffness of 25 in the case of SARS-CoV-2 M^pro^, which we note is about a factor of 2 smaller than the value found by Ming and Wall, suggesting that the optimisation may be structure-dependent.

At first sight, this finding would appear problematic from the point of view of predicting dynamics correlations and allosteric control using ENM models. however, when the thermodynamic quantity of allosteric free energy is calculated for the same range of backbone stiffness (as shown in figure 2d) there is essentially no dependence beyond a slight change developed at a spring stiffness ratio of 10. Although the mode distribution is strongly affected by this enhanced family of models beyond the standard, fixed-stiffness ENM, both fluctuation correlations and the set of key thermodynamic quantities are insensitive, at least for the SARS-CoV-2 M^pro^ homodimer. For this reason, and so that the simulations of the other SARS-CoV-2 proteins of this study may be directly compared to the previous studies on SARS-CoV-2 M^pro^, in the analyses of the following section we employ the standard ENM, with a single bond stiffness between C_*α*_ atoms, ENM node mass equal to 1 atm: while the suitable cut-off was determined by finding the best B-factor correlation with ENM normal modes. The cut-offs used in this study can be found in table 1.

**Table 1.**
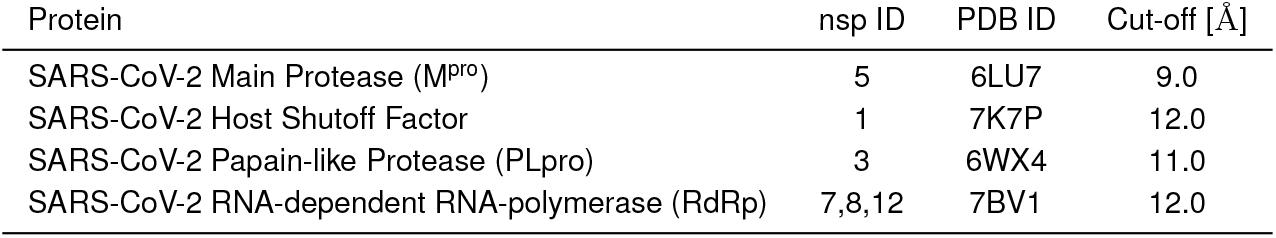
SARS-CoV-2 non-structural proteins’ key information used in this study: non-structural protein ID; PDB ID; ENM cut-off distance based on B-factor correlation.

## 3. Results on three new SARS-CoV-2 proteins

The standard ENM, together with the optimisation of bond-distance and mode-sum cut-offs described in the previous section were applied to three further members of the SARS-CoV-2 protein family. The aims of the computational modelling are twofold: (i) as in previous work on SARS-CoV-2 M^pro^, forensic investigation of the coarse-grained dynamic structure of the proteins may identify accessible residues, sensitive to the control of protein function, that offer the promise of binding targets in future therapeutics; (ii) although there are naturally universal features, each structure is as individual in its dynamic, as in its static, structure. Each case illuminates the physics of protein dynamics, and how it supports the thermodynamics of allosteric control. In particular we shall be interested in the connection between the unperturbed dynamic correlation structure, and the protein response to mutation. Specific pairs of sites whose mutation effects add non-linearly will be of especial interest, as this feature corresponds to the potential for allostery. In each of the three cases below, therefore, we will present data on residue-residue dynamic correlation (compared with a residue proximity map), a spatial structure in which correlations with a particular chosen residue are presented, and maps of two-mutation effects on an appropriate free energy. The latter calculation includes a map of non-linear effects, in which the effect of linear addition of both mutations is subtracted from the combined effect.

### A. Host shutoff factor (nsp1)

In cells infected by SARS-CoV-2, the single-domain protein nsp1 (see figure 3b for a ribbon structure diagram) is implicated centrally in a complex of mechanisms that inhibit host gene expression. Its fold is unusual, consisting of a distorted beta-barrel with an associated helix. Its binding to the 40S ribosomal subunit inhibits host ribosome assembly and also induces endonucleolytic cleavage and degradation of host mRNAs. Of especial interest are therefore residues with strong dynamical correlation with the binding site, which probably implicates, among others, the charged residue K49 (21). Figure 3a reports the strength and sign of residue-residue dynamic correlations (lower right half) and a proximity map (upper left half) for comparison. As in the previous case of SARS-CoV-2 M^pro^, most of the strong correlations arise from spatial proximity, but the contact and correlation maps differ strongly from that previous study, as the nsp1 does not possess the subunit structure of SARS-CoV-2 M^pro^, and both the helix and barrel structure give rise to multiple correlations perpendicular to the main diagonal of the plots. There are three notable regions of correlation that do not correspond to spatial proximity: between residues in the vicinities of 20 and 75, of 30 and 110, and of 45 and 90. The last of these is shown together with the real-space dynamic correlation to residue Q46 in figure 3b. This visualises how the motion of this site, implicated in functional binding, propagates outward to the beta-sheets and helices on the surface of the protein.

**Fig. 3.**
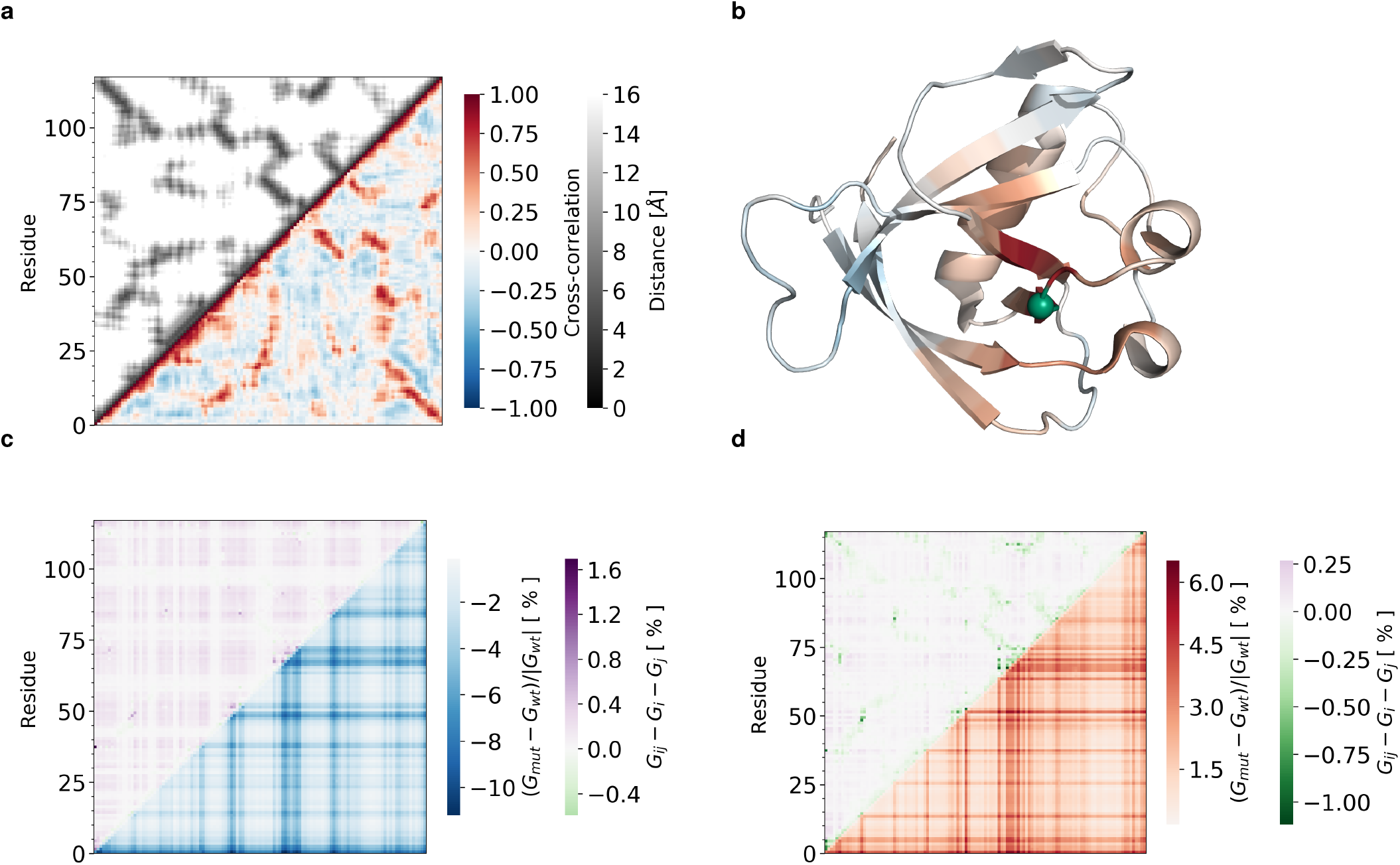
Dynamics of the SARS-CoV-2 Host Shutoff Factor ENM (PDB: 7K7P). **a**, The cross-correlation maps calculated for the first real 25 modes (bottom-right region of the plot) and spacing between residues (top-left region of the plot) for apo form. **b**, A real-space representation of the correlations in 7K7P ENM with respect to residue Q46 (green sphere). The cross correlation matrix was calculated using only the C_*α*_ atoms for the protein. **c**, 2-point mutational map for 7K7P ENM with all possible pairwise combinations of residue mutations with equal spring constant change *k*_*R*_*/k* = 0.25 over the first real 25 fluctuation modes. A map for the fluctuation free energy change (bottom-right region of the plot) plots the relative change in free energy to the wild type ((*G*_*mut*_ − *G*_*wt*_)*/*|*G*_*wt*_|) due to the dimensionless change in the spring constant (*k*_*R*_*/k*) for the mutated residue with the residue number shown. White corresponds to values of free energy predicted by the wild-type ENM. Red corresponds to an increase in (*G*_*mut*_ − *G*_*wt*_)*/*|*G*_*wt*_| (decreased value of *G*_*mut*_ comparing to *G*_*wt*_), whereas blue corresponds to a decrease in ((*G*_*mut*_ − *G*_*wt*_)*/*|*G*_*wt*_|) (increased value of *G*_*mut*_ comparing to *G*_*wt*_). A map for the relative fluctuation free energy change 2-point additivity (top-left region of the plot) plots the difference in the relative change in free energy for 2-point mutation and two single 1-point mutations (*G*_*ij*_ − *G*_*i*_ − *G*_*j*_). White corresponds to *G*_*ij*_ − *G*_*i*_ − *G*_*j*_ = 0.0, i.e. non-synergistic 2-point mutation. Purple corresponds to an increase in *G*_*ij*_ − *G*_*i*_ − *G*_*j*_ (2-point mutation effect on free-energy is greater than combination of two separate 1-point mutations), whereas blue corresponds to a decrease in *G*_*ij*_ − *G*_*i*_ − *G*_*j*_ (2-point mutation effect on free-energy is smaller than combination of two separate 1-point mutations). **d**, 2-point mutational map for 7K7P ENM with all possible pairwise combinations of residue mutations with equal spring constant change *k*_*R*_*/k* = 4.00 over the first real 25 fluctuation modes.

The two-point mutation analyses, by which all springs connected to each of the two residues to relax, or to stiffen, their environment, are displayed in figure 3c and 3d respectively. Because nsp1 is monomeric, with a single but as yet uncertain binding site, results for changes to the total (entropic) free energy (rather than binding, or allosteric free energies as in other cases) are given as a function of the double mutation. Strong control regions for the free energy appear in the vicinities of residues 45-50, 65-70 and 80-85. Non-additive effects of the mutations are displayed by subtracting the free energy changes from the sum of the effects of the two corresponding single mutations (upper left half of plots in 3c and 3d. There is an intriguing difference between the cases of relaxation and stiffening. in the former, non-linear effects are mainly anti-cooperative (in the sense of binding) (increased effect on double mutation), and correlate with pairs in which both mutations have strong effects on the free energy. On stiffening, non-linear effects of both signs are in evidence, with anti-cooperative cases arising as with relaxation. However, co-operative nonlinearities are also now in evidence (coloured green in figure 3d) and correlate with the proximity measure (figure 3a upper half). This is the expected non-linearity from a simple, single mode (’allosteron’) model of entropic free energy change on equal bond stiffening - the two different signs displayed by nsp1 illustrate the two mechanisms of single mode, and mode-mixing allostery proposed in (22).

### B. Papain-like protease (nsp3)

The papain-like protease PLpro enzyme is required for processing viral polyproteins to generate a functional replicase complex. It binds to its target on two sites, on each beta-sheet extremity of its two domains (23). As well as the binding domains for its cleavage activity, PLpro possesses an inhibitor binding site proximal to residue 273. Figure 4 displays the two-point dynamic correlation of the ENM structure for nsp3 based on the PDB file 6WX4 (24) in apo form (panel a) and with an inhibitor bound (ligand VIR251) (panel c). The corresponding real-space correlations mapped to residue 273, adjacent to the inhibitor binding site, are displayed in panels b and d. Note that, as in the other proteins surveyed, most strong correlations are due to spatial proximity, but that the real-space representations help to identify moderately strong (anti-) correlated motion between the inhibitor site and the helical and beta-sheet cleavage protein-binding domains. There are non-proximal positive correlations between 0-50 and 200-220 in both apo and holo1 forms (vertical rectangle on figure 4 a and c), and residues 273 and 293 adjacent to the active site correlate with the protein’s C-terminus.

**Fig. 4.**
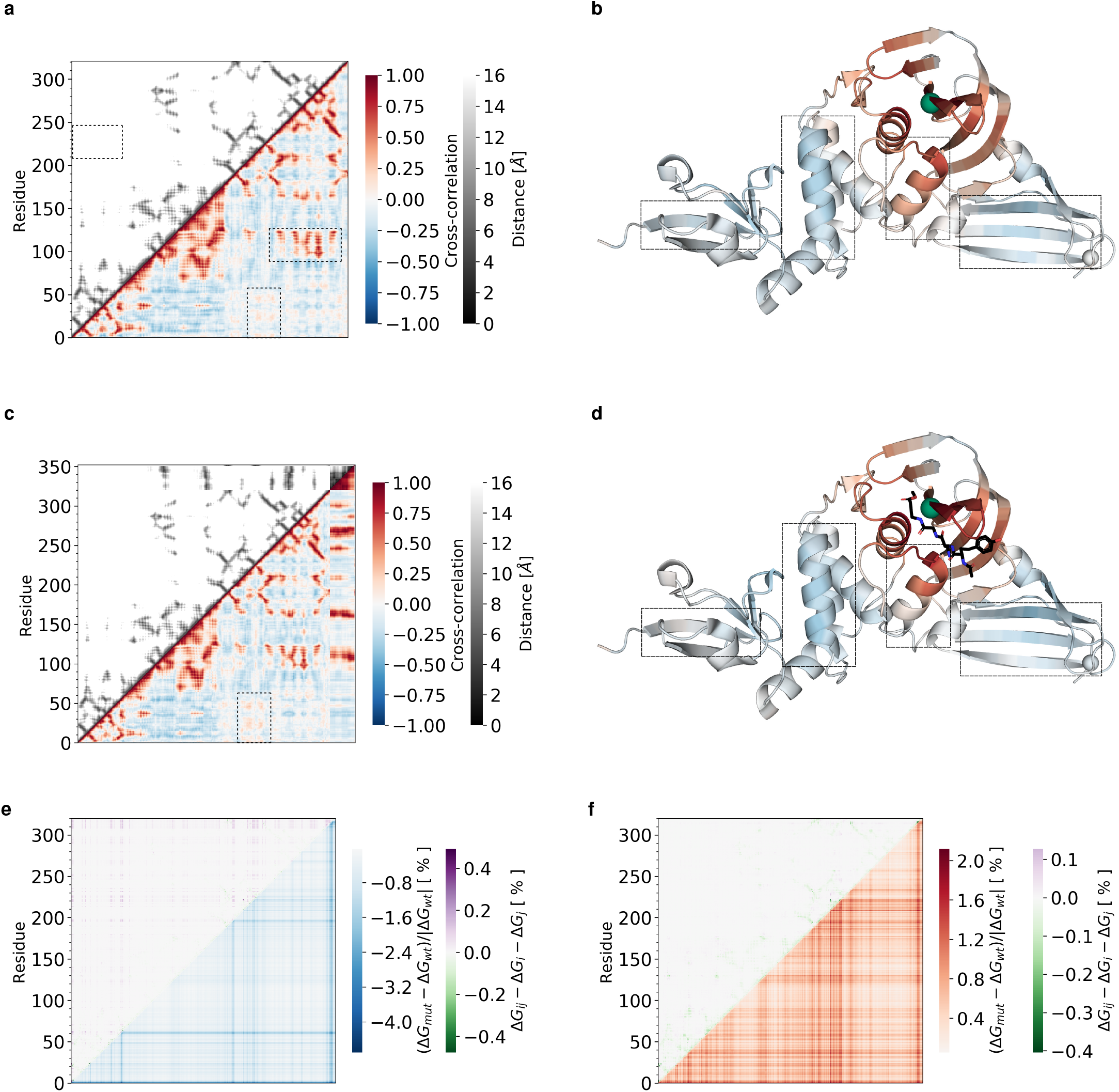
Dynamics of the PLpro ENM dynamics (PDB: 6WX4). **a**, The cross-correlation maps calculated for the first real 25 modes (bottom-right region of the plot) and spacing between residues (top-left region of the plot) for apo form. **b**, A real-space representation of the correlations in 6WX4 ENM with respect to residue 273 (green sphere). The cross correlation matrix was calculated using only the C_*α*_ atoms for the protein. **c**, The cross-correlation maps calculated for the first real 25 modes (bottom-right region of the plot) and spacing between residues (top-left region of the plot) for holo1 form with VIR251 inhibitor. **d**, A real-space representation of the correlations in 6WX4 ENM with respect to residue 273 (green sphere). The cross correlation matrix was calculated using only the C_*α*_ atoms for the protein and all heavy atoms for the ligand (VIR251 inhibitor) shown as sticks. **e**, 2-point mutational map for 6WX4 ENM with all possible pairwise combinations of residue mutations with equal spring constant change *k*_*R*_*/k* = 0.25 over the first real 25 fluctuation modes. A map for the fluctuation free energy change (bottom-right region of the plot) plots the relative change in free energy to the wild type ((Δ*G*_*mut*_ − Δ*G*_*wt*_)*/*|Δ*G*_*wt*_|) due to the dimensionless change in the spring constant (*k*_*R*_*/k*) for the mutated residue with the residue number shown. White corresponds to values of free energy predicted by the wild-type ENM. Red corresponds to an increase in (Δ*G*_*mut*_ − Δ*G*_*wt*_)*/*|Δ*G*_*wt*_| (decreased value of *G*_*mut*_ comparing to Δ*G*_*wt*_), whereas blue corresponds to a decrease in ((Δ*G*_*mut*_ − Δ*G*_*wt*_)*/*|Δ*G*_*wt*_|) (increased value of Δ*G*_*mut*_ comparing to Δ*G*_*wt*_). A map for the relative fluctuation free energy change 2-point additivity (top-left region of the plot) plots the difference in the relative change in free energy for 2-point mutation and two single 1-point mutations (Δ*G*_*ij*_ − Δ*G*_*i*_ − Δ*G*_*j*_). White corresponds to Δ*G*_*ij*_ − Δ*G*_*i*_ − Δ*G*_*j*_ = 0.0, i.e. non-synergistic 2-point mutation. Purple corresponds to an increase in Δ*G*_*ij*_ −Δ*G*_*i*_ −Δ*G*_*j*_ (2-point mutation effect on free-energy is greater than combination of two separate 1-point mutations), whereas blue corresponds to a decrease in Δ*G*_*ij*_ −Δ*G*_*i*_ −Δ*G*_*j*_ (2-point mutation effect on free-energy is smaller than combination of two separate 1-point mutations). **f**, 2-point mutational map for 6WX4 ENM with all possible pairwise combinations of residue mutations with equal spring constant change *k*_*R*_*/k* = 4.00 over the first real 25 fluctuation modes.

There are a few specific points to note in the dynamic structure. First, there are, as with SARS-CoV-2 M^pro^, but even more prominently, ‘H-shaped’ regions of strong positive correlation. This pattern has previously been identified as arising from local hinge-like motions (1). Second, there are notable changes in the cross-correlation between apo and holo1 forms of the protein. There is a strong increase in negative correlation between dynamics at the inhibitor site, and with the inter-domain helix (including residue 250; this is marked by the small rectangle on figure 4c and the vertical rectangle on 4 b and d). Conversely, on inhibitor binding there is a decrease of the negative dynamic correlation on the terminal beta-sheet (horizontal rectangle to left of 4 b and d). Additionally, the negative correlation with the other beta sheet moves towards the end of the protein (three beta-strands on the right of 4 b and d). Finally, the middle helix (right short helix marked by a rectangle in 4 b and d) suffers a decrease in its positive cross-correlation on lingand binding; in the holo1 form the correlation is more focused around the ligand.

The double-mutation scans (of figure 4 e and f) in this case are calculated to measure the effect on free energy of binding at the inhibitor site, so now map a free-energy change rather than a total free energy. However, they possess the same properties in relaxation and stiffening as did nsp1, with the same correlation between co-operative (negative sign) free energy of combination with spatial proximity. Under local bond weakening by a factor of 0.25, the strongest control sites for the inhibitor binding energy are 1-2, 49, 61-2, 196-8 and 315-317. More potential control sites appear when local bonds are stiffened by a factor 4.0, at 1, 37, 120-30, 209, 220-2, 233, 235 and 314-316. Interestingly, the biologically active residues do not appear strongly on the control map, which may indicate an evolved stability to evolutionary point mutation of this family of viral proteins.

### C. RNA-dependent RNA-polymerase (nsp7, 8 and 12)

The final structure of this investigation is the SARS-CoV-2 RNA-dependent RNA polymerase (RdRp), a large 4-chain proteins of over 1000 residues, which regulates viral replication. RdRp (PDB: 7BV1 (25))consists of three non-structural proteins: 12 (chain A), 8 (chains B and D) and 7 (chain C). This enzyme, a much larger structure than the other three proteins of the study, has notwithstanding been proposed as a potential therapeutic target to inhibit viral infection (26). It is, for example, the target of the drug Remdesivir. Because of the size of the protein, we did not perform 2-point mutation scans for this study, but focused on the dynamic correlation map (see figure 1). Residue numbers on the cross-correlation map correspond to index of amino acids and heteroatoms as they appear in the PDB file, because all four chains have missing amino acid regions. In addition to features noted in the other proteins of our sample, the considerably larger size of the RNA-dependent RNA-polymerase permits dynamic structure to emerge at a larger lengthscale. At a more fine-grained view, a set of richly-structured positive cross-correlation profiles are located around the corresponding distance map regions. Negatively correlated regions also populate the map, especially along chain B and almost exclusively along chain D. This effect might be attributed to the incompleteness of chain D (from the point of view of the PDB file) and its distance from the main protein body nsp12.

The amino acid at 557-th (re-indexed residue 478) position in RdRp has been previously shown to promote fidelity in RNA synthesis (27) and is located in the active site (Fig. 5b, dashed square). This is the residue that we have chosen to represent spatial dynamic correlations, colour-coded in figure 1 b, which indicate a positively correlated core region proximal to the reference residue, complemented by a negatively-correlated periphery, some of which appears in helices at a considerable distance. This large structure provides plenty of opportunities for targeting regions away form the active site on the surface of the protein, like nsp8 chain D.

**Fig. 5.**
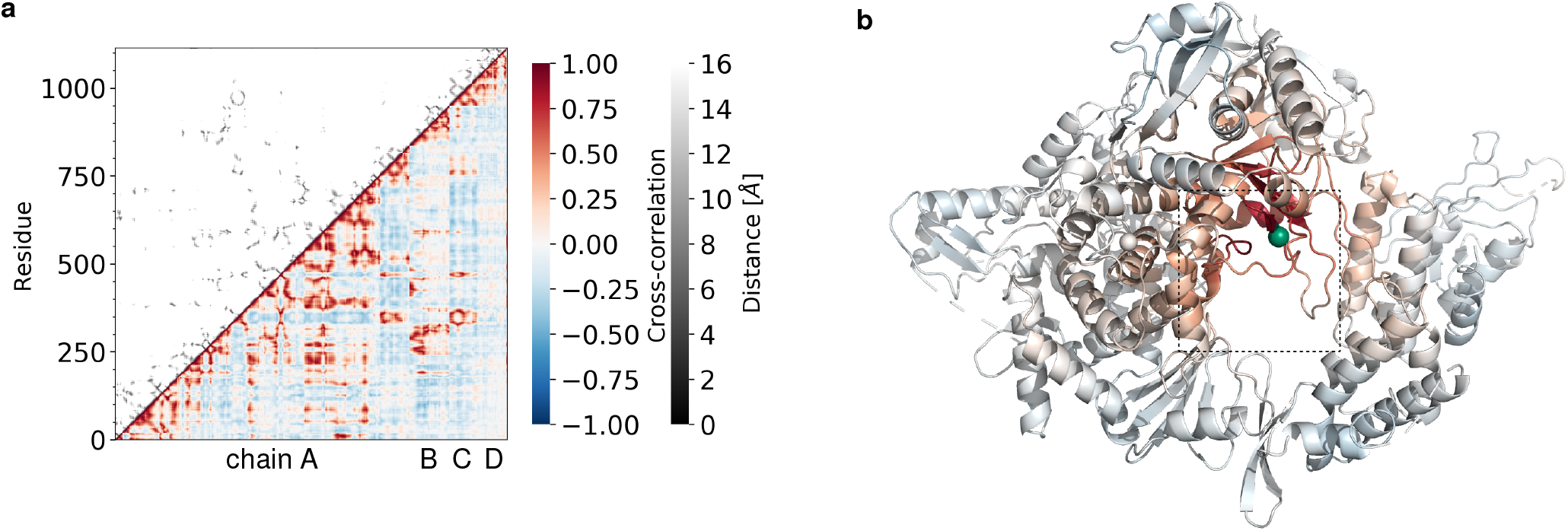
Dynamics of the SARS-CoV-2 Rd-Rp ENM (PDB: 7BV1). **a**, The cross-correlation maps calculated for the first real 25 modes (bottom-right region of the plot) and spacing between residues (top-left region of the plot) for apo form. Labels on x-axis indicate protein chain IDs. **b**, A real-space representation of the correlations in 6WX4 ENM with respect to residue L557 (green sphere), residue 478 on the plot. The cross correlation matrix was calculated using only the C_*α*_ atoms for the protein and heavy heteroatoms, e.g. Zn.

## 4. Discussion

The ENM analysis reinforces previous findings in application to other proteins, including SARS-CoV-2 M^pro^, that local harmonic potentials within the equilibrium protein structure, but without mean structural change can identify candidate biologically active sites for control of protein function in other examples of the SARS-CoV-2 family. Furthermore, it it is not always necessary to possess holo forms of the proteins to locate those active sites, whose correlated, and functional, dynamics are already clear in the apo form. Calculations of those sites where total free energies are sensitive to mutations converge well with the limit of the sum over normal modes. The convergence of calculations of control of the binding or allosteric free energies are more subject to noise, being higher-order difference-quantities, but sufficiently to identify strong candidates for control regions. Computational studies such as this, therefore, accompany and support concurrent experimental programs of scanning for small-molecule binding candidates to the protein.

Because ENM calculations identify entropic contributions to free-energy arising from fluctuations around a mean conformation, they typically deliver a sizeable fraction of the thermal energy *k*_*B*_ *T* per mode affected. This sets the typical scale at a few kJ per mole for the free-energy changes predicted by the model, represented in the results presented here. Previous work has found that experimental free energies are typically ordered in the same sequence as the ENM values, though larger by factors that may arise through coupling of the global modes to unresolved local modes (7, 12, 28).

The ENM models employed in this study was specific for given inhibitors, where these apply. Other ligands might, of course, show different behaviour in the corresponding holo structures and display other “hot-spots”, however, the appearance of active regions, and their coupling, in the apo structure suggests that there are general properties that emerge from the global elastic structure of the protein. The advantage of these coarse-grained studies is that they complement searches for ligands that bind to target proteins by focusing on sites whose local structural dynamics correlate with those at active sites at a lower computational cost than the full molecular MD.

The family of SARS-COV-2 proteins in this study exhibited both universal and specific features in their coarse-grained fluctuation dynamics. In general, positive dynamic residue-residue correlations were dominated by proximity effects, with a few notable allosteric exceptions in each case. On the other hand, strong negative correlations appeared ubiquitously without proximity. New features of coarse-granined dynamics appeared in the largest of the family, the RNA-dependent RNA-polymerase.

As well as providing specific information on its particular proteins, the findings of this study also contribute to the large programme of research on the structure of protein dynamics in general, and on fluctuation-induced allostery without conformational change in particular. As noted above, positive dynamic correlations between distant (allosteric) sites appear to be relatively rare (most strong dynamic correlation arises through spatial proximity between residues). In each case for spatially distant correlations to occur, more than one normal mode anti-nodes of global protein dynamics must coincide.

The study has also identified general ways in which which double mutations contribute in a weakly non-linear addition. Across all proteins of this study, patterns of bond-weakening (through mutation or binding) and bond-strengthening are similar, with the latter exhibiting greater correlation of (negative) non-linearities with spatial proximity. The practically-relevant proteins of this set are therefore able to illustrate examples evolved in nature, of the subtle design principles involved in generating allosteric combinations of either sign, recently theoretically related to the shift of the normal mode structure on the first binding (or bond-stiffening) event (22).

In conclusion, the SARS-COV-2 family of proteins offer a rich set of examples of functional, coarse-grained, protein dynamics. It may be possible to exploit the detailed understanding of these features in future therapeutics; already they have deepened our understanding of how static and dynamic structures of proteins work together in establishing function.

## Materials and Methods

Normal Mode Analysis (NMA) of ENM describes protein motions around equilibrium and can be used to calculate the partition function for large scale harmonic thermal fluctuations in protein structure, including those responsible for allostery (29). Two main approximations of NMA are:

- The structure fluctuates about at local energy minimum. Consequently no other structures beyond the given equilibrium can be explored.
- The force field everywhere arises from sums over ENM harmonic

The whole NMA method can be reduced to three steps:

1. Construct mass-weighted Hessian for a system. For a protein ENM the system consists of the co-ordinates of the C-alpha atoms (*N*) for each residue from the corresponding PDB structure.
2. Diagonalise the mass-weighted Hessian to find eigenvectors and eigenvalues of the normal modes.
3. Calculate the partition function (and so free energy) from the product over the normal mode harmonic oscillations.

The diagonalisation of the 3*N* × 3*N* mass-weighted Hessian matrix is written as

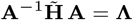

Where 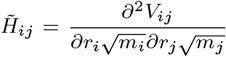: the potential energy function *V*; distance between nodes *r*; node masses *m*. The eigenvectors of the mass-weighted Hessian matrix, columns of **A**, are the normal mode eigenvectors **a**.

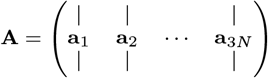

**Λ** is a 3*N* × 3*N* diagonal matrix with diagonal values equal to the associated normal modes’ squared angular frequencies *ω*^2^. The potential function used in this study is:

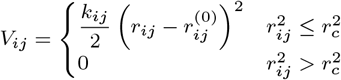

where *r*_*c*_ is a cut-off radius, which for this work is set at 8Å; while *r*^(0)^ is the equilibrium distance between nodes derived form PDB crystallographic structure. For the wild-type protein, all spring constants are equal *k*_*ij*_ = *k*=1 kcalÅ^−2^ mol^−1^.

### Cross-correlation of Motion

The cross-correlation, *C*, is estimated between an ENM node pair as a normalised dot product sum between their normal mode eigenvectors over *v* modes.

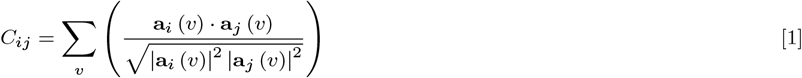

*C* value of 1 implies perfectly correlated motion, −1 perfectly anti-correlated motion and 0 implies totally non-correlated motion.

### Normal Mode Fluctuation Free Energy

Using statistical mechanics it is possible to calculate an estimate to the fluctuation free energy of a system using the frequency of vibrations such as the normal modes. For this method, the partition function for the quantum harmonic oscillator (30), *Z*, for normal mode *k* is given as

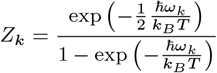

where *k*_*B*_ is the Boltzmann’s constant, 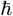 is the reduced Planck’s constant, *T* is temperature in Kelvin and *ω* is, already mentioned, angular frequency. Gibbs free energy (for a given mode) expressed in terms of partition function, with an approximation of little change in volume, can be written as

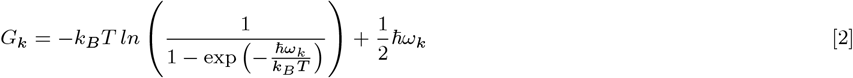

### Ligand Dissociation Constant

When free energy change Δ*G* (**SI, Sec. C**) is known for a dissociation reaction, corresponding dissociation constant *K* can be estimated via

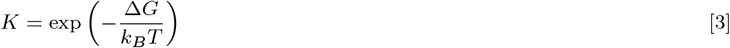

### Molecular Dynamics Simulation

Comparison between Backbone Enhanced ENM, ENM and Molecular Dynamics simulation was carried out by calculating the distributions of eigenfrequencies from the Normal Mode Analysis (NMA) of these three models. The eigenfrequencies for the two ENM models were calculated using the DDPT software while the molecular dynamics eigenfrequencies were calculated by generating a trajectory from the molecular dynamics simulation software Amber (20) and then using DDPT to find the eigenfrequencies from the trajectory. Amber relies on force fields (the Amber force field ff14SB was used (31) in this calculation) to calculate forces between atoms and uses integration algorithms to calculate atom’s trajectories over time. Before a simulation is run, the input protein structure was energy minimised and equilibrated. The pmemd.cuda program was used to implement both pre simulation energy minimisation equilibrations and the production simulation from which the eigenfrequencies are found. CUDA is used to parallelise calculation of the trajectories to run calculations on gpus rather than cpus. The simulations were carried out in an implicit solvent. The amber analysis program cpptraj is used to convert the binary output of pmemed.cuda to a readable trajectory format. The final trajectory from the production simulation is then used to calculate a covariance matrix using the COVAR program in DDPT. COVAR calculates a mass weighted Hessian matrix which is diagonalised. The eigenfrequencies are found through the FREQEN DDPT program.

## ACKNOWLEDGMENTS

ID is grateful for computational support from Research Coding Club at the University of York. TCBM acknowledges support from the EPSRC (UK) through an Established Career Fellowship in the Physics of Life programme. We are grateful to Dr. Alice von der Heydt, Prof. Peter O’Brian and Dr. Sarah Harris for useful discussions. CH would like to acknowledge guidance from Dr. Agnes Noy and George Watson for their guidance and expertise in running the Amber simulations.

